# CardiOvascular examination in awake Orangutans (*Pongo pygmaeus pygmaeus*): Low-stress Echocardiography including Speckle Tracking imaging (the COOLEST method)

**DOI:** 10.1101/2021.07.23.453539

**Authors:** Valérie Chetboul, Didier Concordet, Renaud Tissier, Irène Vonfeld, Camille Poissonnier, Maria Paz Alvarado, Peggy Passavin, Mathilde Gluntz, Solène Lefort, Aude Bourgeois, Dylan Duby, Christelle Hano, Norin Chai

## Abstract

**Introduction:** Cardiovascular diseases have been identified as a major cause of mortality and morbidity in Borneo orangutans (*Pongo pygmaeus pygmaeus*). Transthoracic echocardiography is usually performed under anesthesia in great apes, which may be stressful and risky in cardiac animals. The aim of the present pilot study was hence to develop a quick and non-stressful echocardiographic method (i.e., the COOLEST method) in awake Borneo orangutans (CardiOvascular examination in awake Orangutans: Low-stress Echocardiography including Speckle Tracking imaging) and assess the variability of corresponding variables.

**Materials and Methods:** Four adult Borneo orangutans trained to present their chest to the trainers were involved. A total of 96 TTE examinations were performed on 4 different days by a trained observer examining each orangutans 6 times per day. Each examination included four two-dimensional views, with offline assessment of 28 variables (i.e., two-dimensional (n=12), M-mode and anatomic M-mode (n=6), Doppler (n=7), and speckle tracking imaging (n=3)), representing a total of 2,688 measurements. A general linear model was used to determine the within-day and between-day coefficients of variation.

**Results:** Mean±SD (minimum-maximum) images acquisition duration was 3.8±1.6 minutes (1.3-6.3). All within-day and between-day coefficients of variation but one (n=55/56, 98%) were <15%, and most (51/56, 91%) were <10% including those of speckle tracking systolic strain variables (2.7% to 5.4%).

**Discussion:** Heart morphology as well as global and regional myocardial function can be assessed in awake orangutans with good to excellent repeatability and reproducibility.

**Conclusions:** This non-stressful method may be used for longitudinal cardiac follow-up in awake orangutans.

## Introduction

Cardiovascular diseases have been identified as a major cause of mortality and morbidity in great apes including Borneo orangutans (BO, *Pongo pygmaeus pygmaeus*) under managed care [1]. According to a study based on the North American Orangutan Species Survival Plan, up to 20% of adult orangutan deaths housed in United States and Canadian zoos and private collections between January 1980 and March 2008 were of cardiovascular origin [1]. Based on published reports and analysis of Species Survival Plan necropsy data, orangutans are known to be affected by miscellaneous heart diseases including congenital cardiac defects [2,3], cardiomyopathies [1,4], and viral myocarditis [1,5,6]. The most common heart disease is known as fibrosing cardiomyopathy, characterized by a scattered pattern of myocardial fibrosis with atrophy and hypertrophy of cardiac myocytes, absent or mild inflammation, and no apparent etiology or associated diseases [1].

However, the *antemortem* diagnosis of such heart diseases remains challenging, partly because most of them are responsible for sudden death without prior cardiovascular clinical signs, although congestive heart failure may sometimes develop [1]. Additionally, complementary examinations are considered to be difficult to perform without general anesthesia in these species [1]. Heart diseases in orangutans are therefore currently mainly diagnosed by necropsy and histopathological examinations, and cardiology knowledge in this species remains limited.

Transthoracic echocardiography (TTE) has become a major non-invasive imaging tool for the *antemortem* diagnosis of heart diseases both in humans and various animal species, allowing qualitative description of cardiac abnormalities and quantitative assessment of heart anatomy and function using combined two-dimensional (2D) mode, M-mode, and conventional Doppler modes. More recent advances in ultrasound technology with the introduction of specific imaging modalities, such as 2D speckle tracking imaging (STI), have provided several complementary parameters to assess global and regional myocardial performance [7]. Regarding great apes, a program called the Great Ape Heart Project (GAHP), based at Zoo Atlanta, has collected a database of echocardiograms and other medical information relating to their cardiac health status, and has also provided relevant guidelines for the echocardiographic assessment of anesthetized great apes, including standardization of nomenclature, imaging techniques (M-mode, 2D mode, spectral and color flow Doppler modes), as well as echocardiographic measurements [8]. According to the GAHP report, the use of general anesthesia allows to perform thorough and reliable TTE examinations [8]. However, anesthetizing great apes may be stressful and risky in cardiac animals, which limits the frequency of follow-up TTE sessions. Furthermore, anesthetic drugs may influence some TTE variables [8].

The aims of the present pilot prospective study were therefore 1) to develop and assess the feasibility of a quick and non-stressful TTE method (i.e., the COOLEST method) in awake BO (CardiOvascular examination in awake Orangutans: Low-stress Echocardiography including Speckle Tracking imaging), and then 2) to determine the intra-observer within-day (repeatability) and between-day (reproducibility) variability of the corresponding measurements. To the best of our knowledge, global and regional function of the left ventricle (LV) has never been quantitatively evaluated using STI in this species.

## Materials and Methods

### Animals

Four apparently healthy adult BO were involved in the study, i.e., 3 females (14, 31 and 51 years old; 47, 74 and 69 kg respectively) and 1 male (12 years old; 55 kg). This study was approved by the Ethics Committee of the National Museum of Natural History.

### Transthoracic echocardiographic technique and measurements

#### Echocardiographic procedure

Transthoracic echocardiographic examinations were performed in the orangutans’ indoor enclosure in the Ménagerie, le Zoo du Jardin des Plantes (Paris, France) using a portable cardiovascular ultrasound system (Vivid i, GE Healthcare, 9900 Innovation Drive, Wauwatosa, WI 53226, USA) equipped with a 3S phased-array transducer (1.5-3.5 MHz).

Animal training for the TTE procedure was based on operant conditioning with exclusively positive reinforcement. Orangutans were first trained to sit and present their chest to the trainers against the enclosure mesh with their arms up. They were then desensitized to the ultrasound coupling gel that was generously applied on the thorax through the enclosure mesh, and thereafter to the probe placement on their thorax. After three months of training, the four BO were easily conditioned to stay in the optimal position during several minutes for acquisition of images and video loops.

Transthoracic echocardiograms and corresponding measurements were performed by the same observer (VC), diplomate of the European College of Veterinary Internal Medicine (Cardiology), who was trained to the procedure during five sessions of trial before the beginning of the study.

The final TTE method was the following: trainers were always responsible for bringing BO to an appropriate place, the closest as possible to both the enclosure mesh and the observer. While they were distracting BO by gently and quietly talking to them and giving them some food, optimal acoustic windows were identified and obtained as promptly as possible by the observer, with the transducer placed through the enclosure mesh on BO thorax previously covered by coupling gel (**Vidéo 1; Fig. 1**). Each examination was timed and systematically included four 2D views [8], i.e., the apical 4- and 5-chamber views, the parasternal short-axis view at the level of the aortic valve, and the apical long-axis view optimized for tricuspid annular plane systolic excursion (TAPSE) measurement.

**Fig 1:**
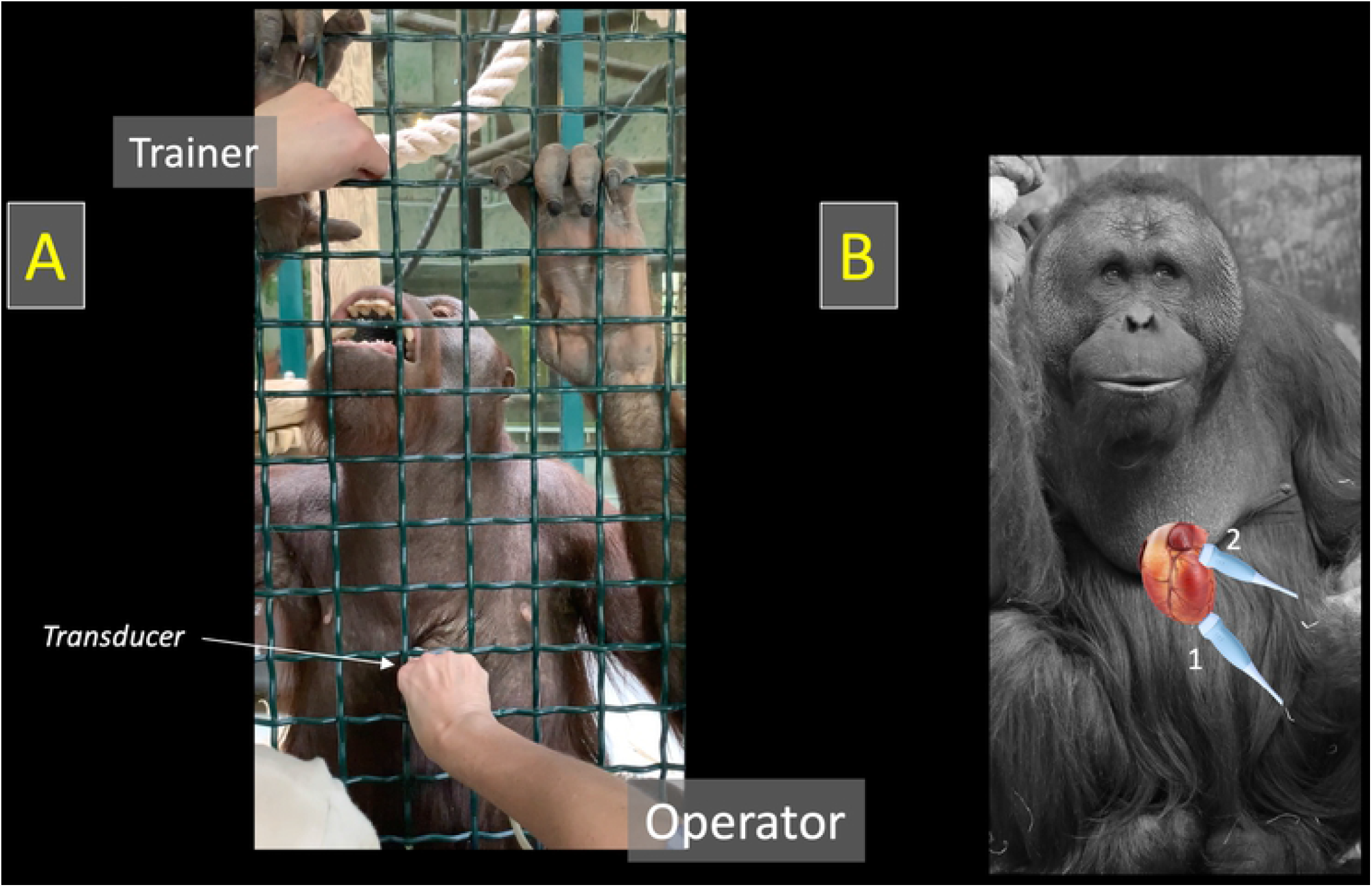
Orangutan positioning for echocardiographic examination and probe placement. Orangutan positioning for echocardiographic examination (A) and probe placement (B) for obtaining the apical long-axis views (1) and the parasternal short-axis view at the level of the aortic valve (2). *Figure 1B : courtesy of Emmanuel Baril (Ménagerie, le zoo du jardin de plantes, Paris)*.

**Video 1: Video showing one of the Borneo orangutans (Pongo pygmaeus pygmaeus) involved in the study**. The animal is sitting and presenting its chest to the trainer against the enclosure mesh with its arms raised, for recording transthoracic echocardiographic images and video loops. The transducer is placed by the observer through the enclosure mesh, after generously applying coupling gel to the acoustic window, while the trainer is talking and feeding the orangutan.

#### Apical 4-chamber-view (Figs. 2 and 3)

The apical 4-chamber was obtained as described in the GAHP report, with the probe placed on the left lateral portion of the chest at the apex of the heart [8].

M-mode echocardiograms showing at least five consecutive cardiac cycles were obtained for offline mitral annular motion measurement (**Fig. 2A**) [9]. Pulsed-wave Doppler images were also obtained for offline assessment of peak velocity of early and late diastolic transmitral flows (E and A waves, respectively) and E wave deceleration time, with calculation of the E:A ratio (**Fig. 2B**). Heart rate was assessed by calculating the time interval between two peak velocities of early diastolic transmitral flow.

**Fig 2:**
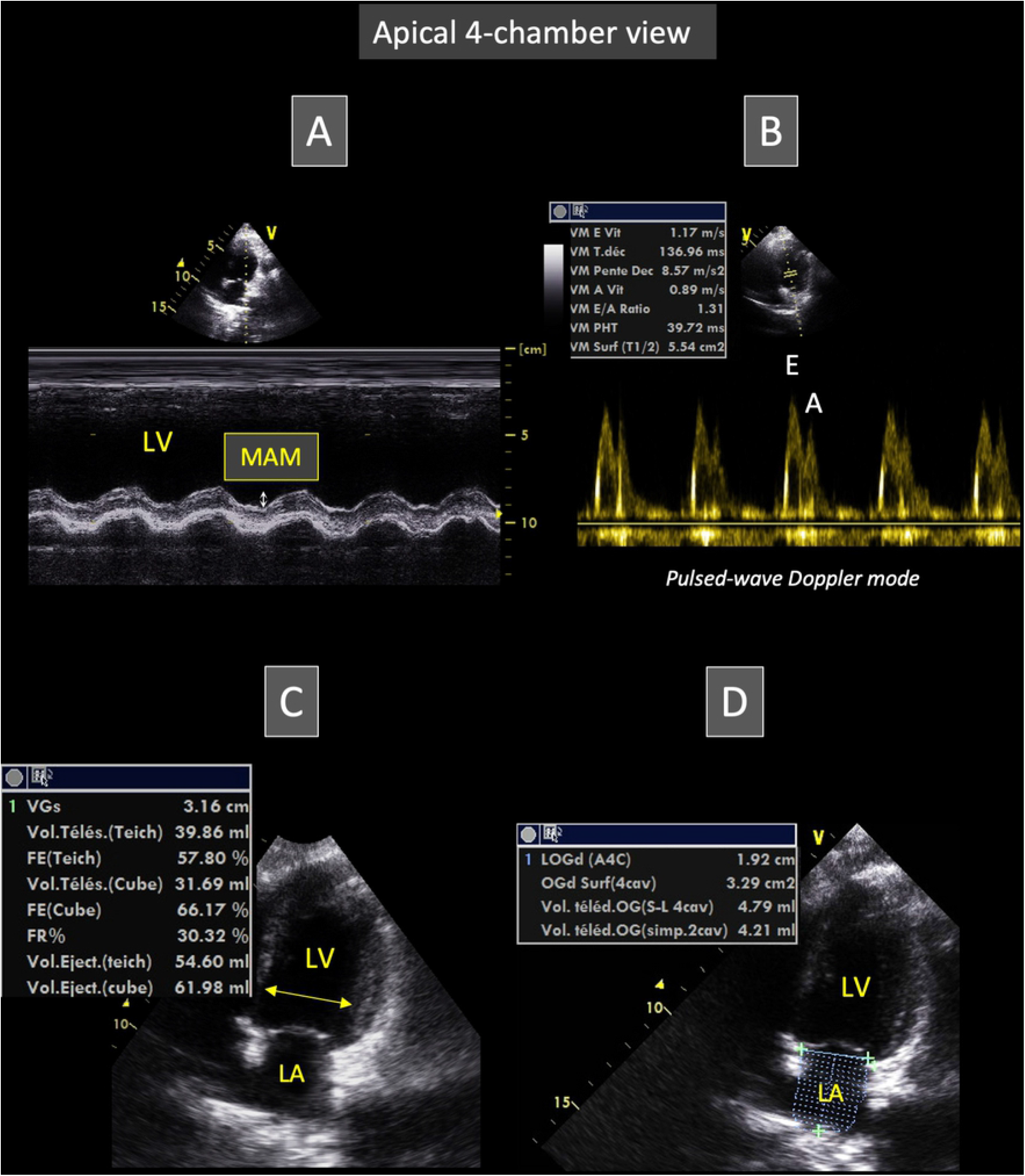
Representative apical 4-chamber view obtained from Borneo orangutans involved in the study. This view allows to place the M-mode cursor through the lateral mitral annulus for measurement of mitral annular motion (MAM, A) and to obtain transmitral flow recordings using pulsed-wave Doppler mode (B). Video loops of this view can also be used for left ventricular (C) and left atrial measurements (D). Figure 2C and Figure 2D respectively show measurements of the end-systolic left ventricular diameter and end-diastolic left atrial volume (see text for explanation). *E and A: peak velocity of early and late diastolic transmitral flows, respectively; LA: left atrium; LV: left ventricle*.

The apical 4-chamber view was also recorded as video loops including at least five consecutive cardiac cycles, that were digitally stored for offline analysis and assessment of left heart dimensions and function. The end-diastolic and end-systolic LV internal diameters were measured from inner edge to inner edge of the LV cavity (**Fig. 2C**), perpendicular to myocardial walls, and just proximal to mitral leaflet tips at their maximal diastolic opening. End-diastole was defined as the frame showing the largest LV cavity after mitral valve closure, and end-systole as the last frame before mitral valve opening. The corresponding fractional shortening was then calculated. Additionally, the end-diastolic and end-systolic LV volumes were automatically assessed using the Simpson’s method of discs by manually tracing the LV endocardial border at blood-tissue interface, from the septal mitral annulus, up to the apex, and then to the lateral mitral annulus, with exclusion of papillary muscles from the traced border [10]. Ventricular volumes were then used to calculate the LV ejection fraction as previously described [10]. From the same 2D loops of the apical 4-chamber view used to assess LV dimensions, the endocardial border of the left atrium (LA) was manually traced at end-systole and end-diastole, with exclusion of the left auricle and pulmonary veins from the traced border, to assess LA lengths and volumes using the Simpson’s method of discs (**Fig. 2D**) [11]. Left atrial volumes were then indexed to body weight.

Furthermore, maximal thickness of the anterior and posterior mitral valve leaflets was measured from the same 2D loops at their maximal diastolic opening.

Finally, the apical 4-chamber loops were also used to assess the STI longitudinal LV systolic strain (StS), as previously described (**Fig. 3**) [12]. Briefly, the LV endocardial border was manually traced at end-diastole (corresponding to the largest LV cavity after mitral valve closure). A region of interest, in which the computer software (Echo Pac PC 6.3 software for Vivid 7, GE Healthcare) automatically performed speckle tracking, was drawn to include the entire myocardium, and the software algorithm then automatically segmented the LV myocardium into 6 segments (3 within the interventricular septum and 3 within the LV free wall). Six longitudinal StS profiles were thus obtained, and peak longitudinal StS values were automatically assessed for each of the 6 LV segments. The mean value of peak StS (corresponding to StS_4chv_) was then calculated.

**Fig 3:**
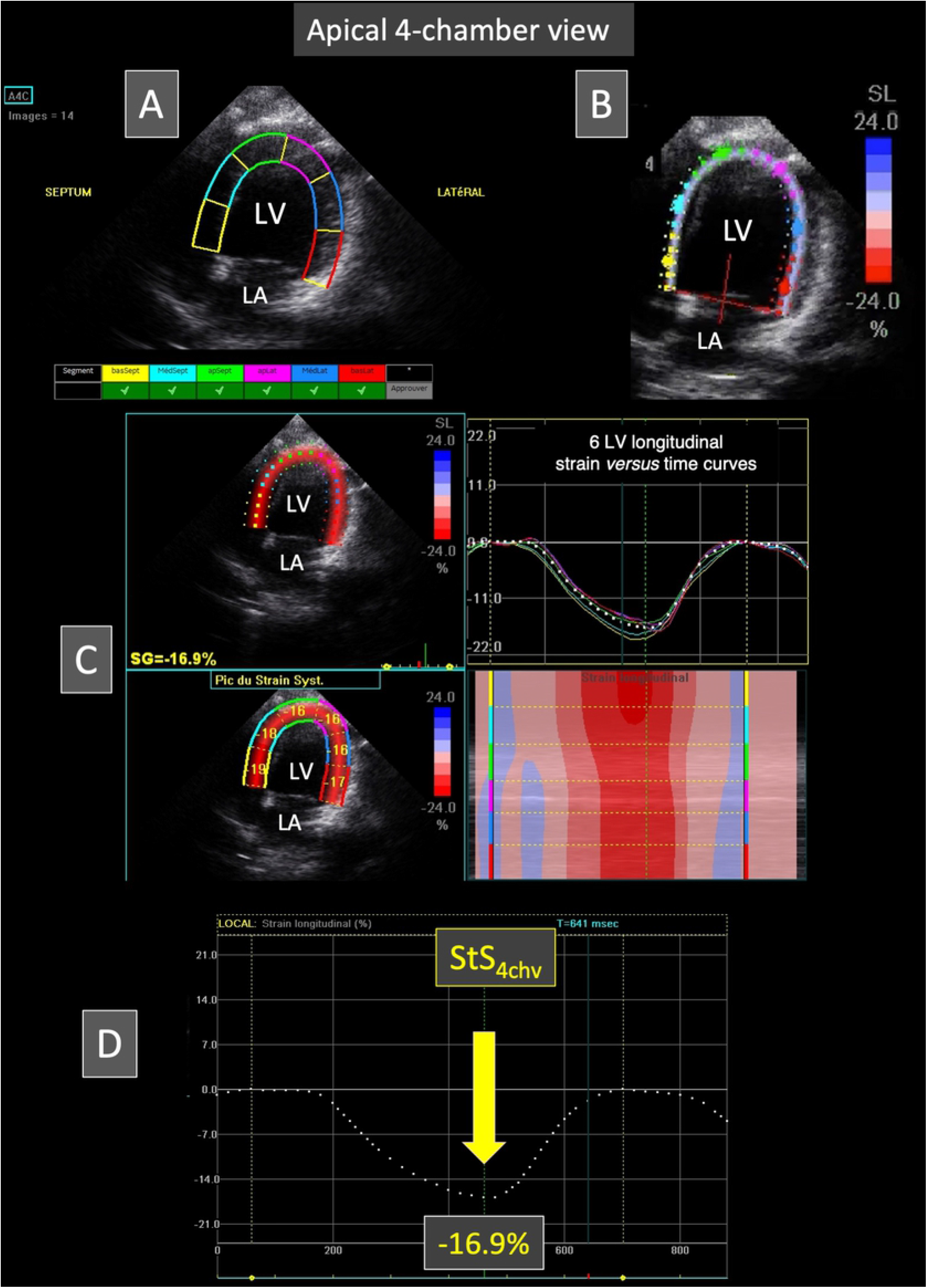
Representative speckle tracking imaging examination performed from the apical 4-chamber view in a Borneo orangutan. The left ventricular endocardial border is traced and a region of interest including left ventricular myocardial walls is automatically drawn (A). The software algorithm then automatically separates left ventricular myocardial walls into 6 equidistant segments (A and B). The tracking quality is displayed for each segment (A). Figure C shows on the right the 6 left ventricular longitudinal strain *versus* time curves (corresponding to the 6 myocardial segments) during a single cardiac cycle. This representative case demonstrates that all the 6 LV segments undergo a relatively homogenous systolic myocardial shortening during systole (negative strain). The corresponding color map below displays the change in strain over time in each segment during the same single cardiac cycle. The peak systolic strain is displayed in each of the six segments on the two-dimensional color-coded view (left). Figure D shows the mean left ventricular longitudinal strain *versus* time curve. In this case, the peak systolic strain value (StS_4chv_) is - 16.9%. *LA: left atrium; LV: left ventricle*.

#### Apical 5-chamber-view (Figs. 4 and 5)

The apical 5-chamber view was obtained from the apical 4-chamber view as described in the GAHP report, by tilting and sliding the transducer beam cranially [8]. Continuous-wave and pulsed-wave Doppler images were obtained for offline calculation of aortic peak flow velocity and isovolumic relaxation time, respectively (**Fig. 4A**). Additionally, the apical 5-chamber view was recorded as video loops including at least five consecutive cardiac cycles, that were digitally stored for offline analysis. Anatomic M-mode (AMM) images were generated from these digital 2D video loops. The electronic AMM cursor was carefully placed across the LA and the aortic root, perpendicular to the aortic and LA walls (**Fig. 4B**). The LA end-systolic AMM diameter and the aortic end-diastolic AMM diameter were measured from inner edge to inner edge, and the corresponding LA:Aorta AMM ratio was then calculated.

**Fig 4:**
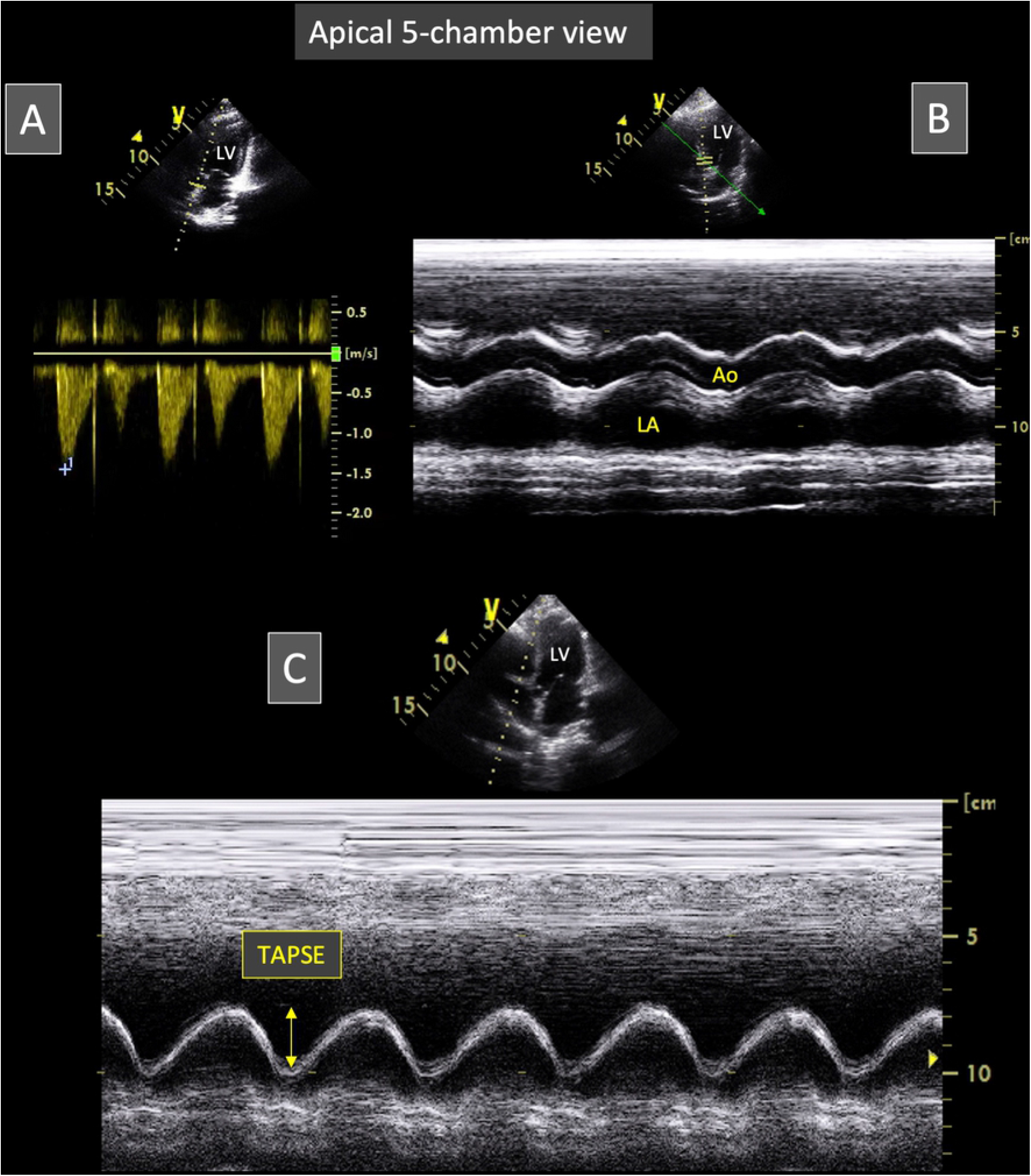
Representative apical 5-chamber view obtained from Borneo orangutans involved in the study. This view allows to obtain aortic flow recordings using continuous-wave Doppler mode (A). Video loops of this view can also be recorded for post-processing measurements of the left atrium and the aorta using the anatomic M-mode technique (B). For this purpose, the M-mode line is positioned perpendicular to left atrial and aortic walls (B). Lastly, by slightly tilting the probe from this position, the M-mode line can also be placed through the lateral segment of the tricuspid annulus for optimal measurement of the tricuspid annular plane systolic excursion (TAPSE; C). *Ao: aorta; LA: left atrium; LV: left ventricle*.

The same video loops were used to assess the STI longitudinal LV StS (StS_5chv_), as described above for StS_4chv_ (**Fig. 5**). The global LV StS (StS_global_) was assessed by calculating the mean of StS_5chv_ and StS_4chv_.

**Fig 5:**
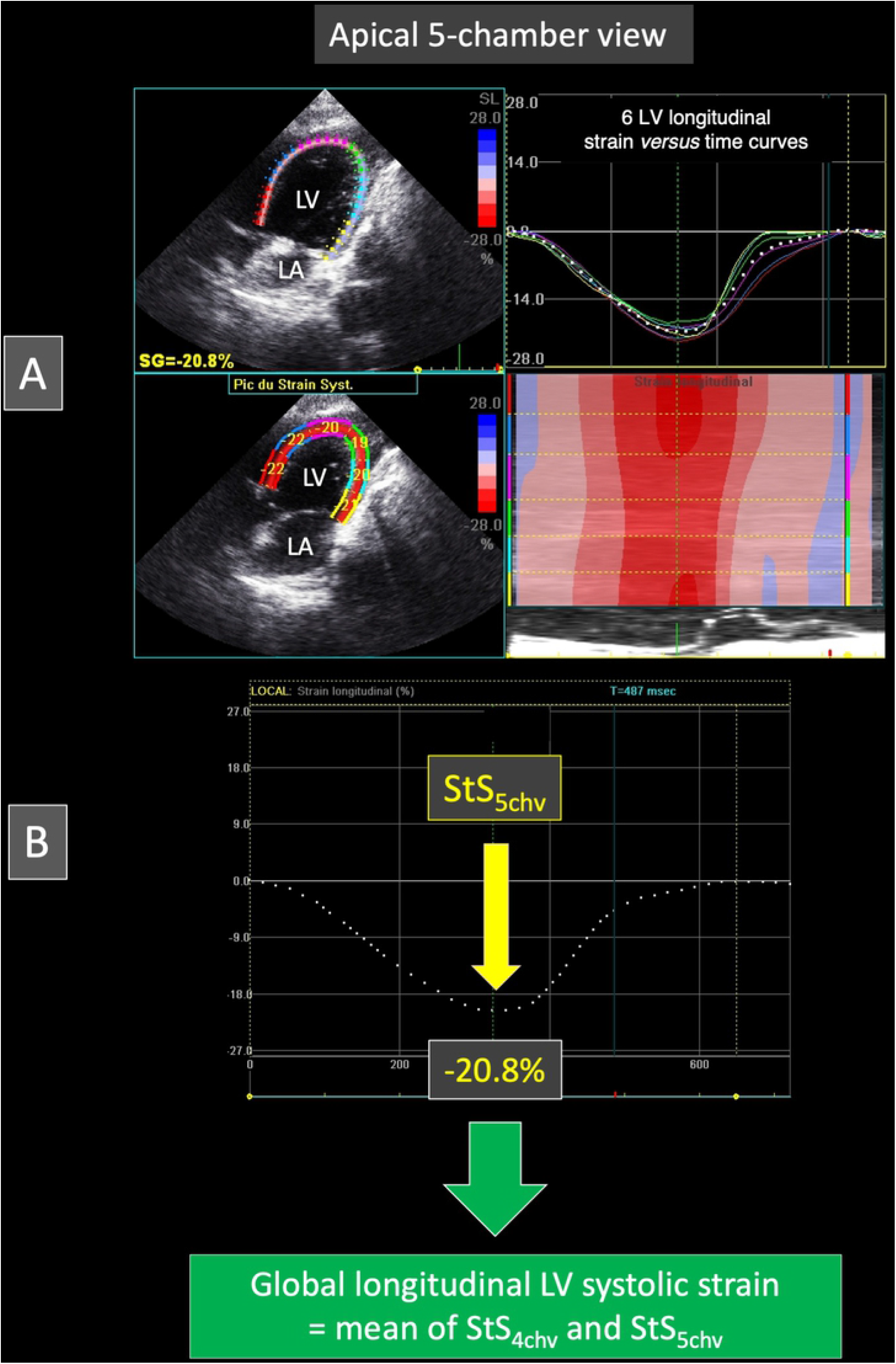
Representative speckle tracking imaging examination performed from the apical 5-chamber view in a Borneo orangutan. As described in Figure 3 for the apical 4-chamber view, the left ventricular endocardial border has been traced and a region of interest including left ventricular myocardial walls has been automatically drawn. Six equidistant myocardial segments have been defined. Figure A shows on the right the 6 left ventricular longitudinal strain *versus* time curves (corresponding to the 6 myocardial segments) during a single cardiac cycle. This representative case demonstrates that all the 6 LV segments undergo a relatively homogenous systolic myocardial shortening during systole (negative strain), with the corresponding color map below displaying the change in strain over time in each segment during the same single cardiac cycle, and with the peak systolic strain displayed in each segment on the two-dimensional color-coded view (left). Figure B shows the mean left ventricular longitudinal strain *versus* time curve. In this case, the peak systolic strain value (StS_5chv_) is - 20.8%. The global left ventricular systolic strain is then assessed by averaging mean peak systolic strain values obtained from the apical 4- and 5-chamber view, i.e., StS_4chv_ and StS_5chv_, respectively. *LA: left atrium; LV: left ventricle*.

Heart rate was assessed by calculating the time interval between two aortic peak flow velocities.

#### Parasternal short-axis view at the level of the aortic valve

The parasternal short-axis view at the level of the aortic valve was obtained as described in the GAHP report, with the probe placed more cranially as compared to the two above-mentioned apical views, between the sternum and the left nipple [8]. Continuous-wave Doppler images were obtained for offline calculation of pulmonic peak flow velocity.

#### Apical long-axis view optimized for TAPSE measurement

M-mode echocardiograms showing at least five consecutive cardiac cycles were obtained using either apical 4- or 5-chamber view (**Fig. 4C**), slightly modified for optimal visualization of the tricuspid annulus and offline TAPSE measurement. In addition, TAPSE values were indexed to body weight. Heart rate was assessed by calculating the time interval between two maximal tricuspid annular systolic excursions on M-mode echocardiograms.

### Assessment of intra-observer within-day and between-day variability of TTE variables

A total of 96 echocardiographic examinations were performed on 4 different days over a 2.5-month period. Each BO was examined 6 times per day. As above-mentioned, each examination performed by the same trained observer included four two-dimensional (2D) views, with offline assessment of 28 variables (i.e., 2D (n=12), M-mode and AMM (n=6), Doppler (n=7), and STI (n=3)), representing a total of 2,688 measurements. A general linear model was used to determine the within-day and between-day coefficients of variation (CV).

### Statistical analysis

Data are expressed as mean ± standard deviation (SD). Statistical analyses were performed using a computer software (R Core Team (2018). R version 3.5.2: A language and environment for statistical computing. R Foundation for Statistical Computing, Vienna, Austria). Briefly, and as already described in our previous ultrasound imaging validation studies performed on wild or exotic animals [13, 14], the following linear model was used to analyze the within-day and between-day variability of the TTE variables:

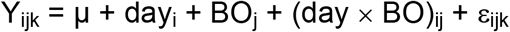

where Y_ijk_ is the k^th^ value measured for BO j on day i, μ is the general mean, day_i_ is the differential effect of day i, BO_j_ is the differential effect of BO j, (day × BO)_ij_ is the interaction term between day and BO, and ε_ijk_ is the model error. The SD of repeatability was estimated as the residual SD of the model and the SD of reproducibility as the SD of the differential effect of day. The corresponding CV values were determined by dividing each SD by the mean. The level of significance was set at P<0.05.

## Results

All 96 TTE examinations could be performed, with a mean image acquisition duration of 3.8 ± 1.6 minutes (1.3-6.3).

All of the 28 tested variables (i.e., 2D (n=12), M-mode and anatomic M-mode (n=6), Doppler (n=7), and STI (n=3)) could be assessed offline, representing a total of 2,688 measurements.

The mean heart rate ± SD during TTE examinations was 93 ± 9 beats per minute (76-118).

**Table 1** and **Table 2** give the values for the 2,688 repeated TTE measurements, i.e., 2D, M-mode and AMM variables in **Table 1**, and Doppler and STI variables in **Table 2**. The corresponding within-day and between-day SD and CV values are shown in **Tables 3 and 4**, respectively.

**Table 1:** Mean ± standard deviation (SD), minimum and maximum values of repeated measurements of two-dimensional (n=12), M-mode (n=3) and anatomic M-mode (n=3) echocardiographic variables (n=18) obtained by a trained observer in 4 Borneo orangutans (*Pongo pygmaeus pygmaeus*) from 96 transthoracic examinations.

| **Echocardiographic parameters** | **Mean ± SD** | **Minimum - Maximum** |
| --- | --- | --- |
| ***Apical 4-chamber view two-dimensional-parameters*** |  |  |
| Left ventricular end-diastolic internal diameter (mm) | 40.0 ± 4.3 | 34.7 – 47.2 |
| Left ventricular end-systolic internal diameter (mm) | 27.4 ± 3.0 | 23.6 – 32.8 |
| Left ventricular radial fractional shortening (%) | 31.5 ± 1.9 | 20.8 – 34.6 |
| Left ventricular ejection fraction (%) | 52.6 ± 8.4 | 40.0 – 69.0 |
| Left atrial end-diastolic length (cm) | 2.09 ± 0.27 | 1.50 – 2.80 |
| Left atrial end-systolic length (cm) | 3.07 ± 0.36 | 2.40 – 3.90 |
| Left atrial end-diastolic volume (mL) | 5.8 ± 1.7 | 3.1 – 8.9 |
| Left atrial end-diastolic volume indexed to body weight (mL/kg) | 0.09 ± 0.02 | 0.06 – 0.13 |
| Left atrial end-systolic volume (mL) | 17.5 ± 3.6 | 13.0 – 23.6 |
| Left atrial end-systolic volume indexed to body weight (mL/kg) | 0.29 ± 0.02 | 0.25 – 0.34 |
| Maximal thickness of the anterior mitral valve leaflet (mm) | 3.2 ± 0.6 | 2.3 – 4.3 |
| Maximal thickness of the posterior mitral valve leaflet (mm) | 3.2 ± 0.7 | 2.1 – 4.4 |
| ***Apical 4-chamber view M-mode parameter*** |  |  |
| Mitral annular motion (cm) | 1.45 ± 0.10 | 1.28 – 1.60 |
| ***Apical 5-chamber view anatomic M-mode parameters*** |  |  |
| Left atrial end-systolic diameter (cm) | 2.83 ± 0.38 | 2.33 – 3.51 |
| Aortic end-diastolic diameter (cm) | 2.28 ± 0.25 | 1.08 – 2.54 |
| Left atrium/aorta ratio | 1.24 ± 0.20 | 1.02 – 2.38 |
| ***Apical long-axis view optimized for M-mode tricuspid annular plane systolic excursion measurement*** |  |  |
| Tricuspid annular plane systolic excursion (cm) | 2.55 ± 0.38 | 1.90 – 3.20 |
| Tricuspid annular plane systolic excursion indexed to body weight (mm/kg) | 0.42 ± 0.02 | 0.38 – 0.47 |

**Table 2:** Mean ± standard deviation (SD), minimum and maximum values of repeated measurements of Doppler (n=7) and STE (n=3) variables (n=10) obtained by a trained observer in 4 Borneo orangutans (*Pongo pygmaeus pygmaeus*) from 96 transthoracic examinations.

| Echocardiographic parameters | Mean ± SD | Minimum - Maximum |
| --- | --- | --- |
| ***Doppler parameters*** |  |  |
| Peak velocity of early diastolic transmitral flow (E wave) (m/s) | 1.19 ± 0.10 | 1.04 – 1.42 |
| Peak velocity of late diastolic transmitral flow (A wave) (m/s) | 0.61 ± 0.20 | 0.30 – 0.94 |
| E:A ratio | 2.21 ± 0.79 | 1.20 – 3.70 |
| E wave deceleration time (ms) | 125.7 ± 13.8 | 101.0 – 166.0 |
| Aortic peak flow velocity (m/s) | 1.52 ± 0.12 | 1.30 – 1.77 |
| Isovolumic relaxation time (ms) | 76.3 ± 17.2 | 50.0 – 104.0 |
| Pulmonic peak flow velocity (m/s) | 1.25 ± 0.10 | 1.07 – 1.45 |
| ***Speckle tracking imaging parameters*** |  |  |
| Longitudinal left ventricular systolic strain (4-chamber view) x -1 (%) | 16.3 ± 0.7 | 14.7 – 17.8 |
| Longitudinal left ventricular systolic strain (5-chamber view) x -1 (%) | 18.1 ± 1.3 | 15.9 – 20.8 |
| Global left ventricular systolic strain x -1 (%) | 17.2 ± 0.6 | 16.1 – 18.4 |

**Table 3:** Within-day and between-day variability, expressed as standard deviations (SD) and coefficients of variation (CV), of two-dimensional (n=12), M-mode (n=3) and anatomic M-mode (n=3) echocardiographic variables (n=18) obtained by a trained observer in 4 Borneo orangutans (*Pongo pygmaeus pygmaeus*) from 96 transthoracic examinations.

| Echocardiographic parameters | Within-day |  | Between-day |  |
| --- | --- | --- | --- | --- |
|  | SD | CV (%) | SD | CV (%) |
| ***Apical 4-chamber view two-dimensional-parameters*** |  |  |  |  |
| Left ventricular end-diastolic internal diameter (mm) | 0.88 | 2.20 | 1.17 | 2.94 |
| Left ventricular end-systolic internal diameter (mm) | 0.70 | 2.57 | 1.18 | 4.30 |
| Left ventricular radial fractional shortening (%) | 1.57 | 4.99 | 1.88 | 5.98 |
| Left ventricular ejection fraction (%) | 2.68 | 5.10 | 3.81 | 7.26 |
| Left atrial end-diastolic length (cm) | 0.18 | 8.43 | 0.35 | 16.78 |
| Left atrial end-systolic length (cm) | 0.19 | 6.10 | 0.28 | 9.07 |
| Left atrial end-diastolic volume (mL) | 0.36 | 6.27 | 0.39 | 6.79 |
| Left atrial end-diastolic volume indexed to body weight (mL/kg) | 0.01 | 6.23 | 0.01 | 7.10 |
| Left atrial end-systolic volume (mL) | 0.78 | 4.48 | 1.27 | 7.25 |
| Left atrial end-systolic volume indexed to body weight (mL/kg) | 0.01 | 4.33 | 0.02 | 7.01 |
| Maximal thickness of the anterior mitral valve leaflet (mm) | 0.16 | 4.81 | 0.24 | 7.34 |
| Maximal thickness of the posterior mitral valve leaflet (mm) | 0.19 | 5.96 | 0.32 | 10.08 |
| ***Apical 4-chamber view M-mode parameter*** |  |  |  |  |
| Mitral annular motion (cm) | 0.05 | 3.66 | 0.08 | 5.75 |
| ***Apical 5-chamber view anatomic M-mode parameters*** |  |  |  |  |
| Left atrial end-systolic diameter (cm) | 0.09 | 3.13 | 0.13 | 4.68 |
| Aortic end-diastolic diameter (cm) | 0.10 | 4.45 | 0.16 | 7.04 |
| Left atrium/aorta ratio | 0.12 | 9.73 | 0.18 | 14.82 |
| ***Apical long-axis view optimized for M-mode tricuspid annular plane systolic excursion measurement*** |  |  |  |  |
| Tricuspid annular plane systolic excursion (cm) | 0.11 | 4.10 | 0.18 | 7.15 |
| Tricuspid annular plane systolic excursion indexed to body weight (mm/kg) | 0.02 | 4.11 | 0.03 | 6.44 |

**Table 4:** Within-day and between-day variability, expressed as standard deviations (SD) and coefficients of variation (CV), of Doppler (n=7) and STE (n=3) variables (n=10) obtained by a trained observer in 4 Borneo orangutans (*Pongo pygmaeus pygmaeus*) from 96 transthoracic examinations.

| Echocardiographic parameters | Within-day |  | Between-day |  |
| --- | --- | --- | --- | --- |
|  | SD | CV (%) | SD | CV (%) |
| ***Doppler parameters*** |  |  |  |  |
| Peak velocity of early diastolic transmitral flow (E wave) (m/s) | 0.03 | 2.46 | 0.04 | 3.29 |
| Peak velocity of late diastolic transmitral flow (A wave) (m/s) | 0.04 | 6.40 | 0.07 | 12.05 |
| E:A ratio | 0.17 | 7.54 | 0.30 | 13.43 |
| E wave deceleration time (ms) | 5.80 | 4.62 | 10.17 | 8.09 |
| Aortic peak flow velocity (m/s) | 0.04 | 2.33 | 0.07 | 4.79 |
| Isovolumic relaxation time (ms) | 2.76 | 3.62 | 3.68 | 4.83 |
| Pulmonic peak flow velocity (m/s) | 0.04 | 3.29 | 0.06 | 4.79 |
| ***Speckle tracking imaging parameters*** |  |  |  |  |
| Longitudinal left ventricular systolic strain (4-chamber view) x -1 (%) | 0.66 | 4.04 | 0.89 | 5.44 |
| Longitudinal left ventricular systolic strain (5-chamber view) x -1 (%) | 0.67 | 3.70 | 0.87 | 4.82 |
| Global longitudinal left ventricular systolic strain (%) | 0.46 | 2.68 | 0.73 | 4.26 |

Nearly all within-day and between-day CV values (55/56, 98%) were <15%, except for the between-day CV value of the end-diastolic left atrial length (16.8%).

Most within-day and between-day CV values (51/56, 91%) were <10%, including those of STI StS variables (2.7% to 5.4%), the lowest being observed for the end-diastolic LV internal diameter (2.2% and 2.9% for within- and between-day CV values, respectively).

A significant BO effect was observed for all TTE variables (P<0.05), except for the LV fractional shortening (P=0.33). No significant day effect was observed for all of the 28 tested variables, except for the less reproductible one, i.e., the end-diastolic left atrial length (P=0.04), and for the peak aortic flow velocity (P=0.03).

## Discussion

Knowledge of cardiology in BO is scarce. To the best of the authors’ knowledge, no study has yet provided quantitative data on heart morphology and function in awake BO, and LV myocardial function has never been quantitatively evaluated using STI in great apes. As optimal diagnostic techniques require precision (e.g., repeatability and reproducibility), accuracy, specificity, and sensitivity, the present study has investigated the feasibility and variability of TTE including STI in awake BO. This non-invasive and non-stressful imaging approach represents an essential initial step in the development of valuable diagnostic tools for the non-invasive *antemortem* diagnosis and longitudinal follow-up of BO heart diseases.

Borneo orangutans are classified as critically endangered according to the International Union for the Conservation of Nature [15]. Main causes of BO population decline include habitat destruction (deforestation, bushfire), illegal hunting, climate change, and lack of awareness of local populations [15]. As a result, captive BO are of major importance to increase public awareness and raise funds for their conservation, and also to act as a potential source of repopulation [4]. Captive BO population management is hence essential to maintain healthy individuals under managed care, and in this context any disease can be considered as a threat [4]. Cardiovascular diseases are a major cause of morbidity and mortality in all great ape taxa, accounting for up to 20%, 38%, 41%, and 45% of deaths in captive orangutans (*Pongo spp*.), chimpanzees (*Pan troglotydes*), gorillas (*Gorilla spp*.) and bonobos (*Pan paniscus*), respectively [1]. Transthoracic echocardiography could therefore represent a useful non-invasive tool for early diagnosis and routine follow-ups of great ape heart diseases. Additionally, TTE could allow a better understanding of these pathological conditions, including their etiology and pathogenesis, which would be key to the conservation of these species.

As stated in the GAHP report, performing TTE under general anesthesia provides better-quality images, allows thorough examinations, and also offers the possibility of examining other systems in addition to the cardiovascular one [8]. However, despite these benefits, anesthesia may be stressful and risky in cardiac animals, which limits the use of TTE as a routine diagnostic tool. Moreover, health issues, namely respiratory infectious diseases, appear to be of specific concern in BO [16], which may also increase anesthetic risks. This is the reason why the present report was specifically dedicated to TTE in awake BO.

The innovative veterinary program called the GAHP has provided guidelines for TTE in great apes, with suggestions for standardization of nomenclature, cardiac imaging views, and measurements [8]. Several views have thus been identified and described, i.e., three parasternal long axis views, three parasternal short-axis views, three apical views (apical 5-, 4- and 2-chamber views), and also one suprasternal and one subcostal view. The objective of the present study was to develop a TTE method in awake BO, that should ideally be non-stressful for the animals and as safe as possible for involved observers, trainers, and equipment. Therefore, time for image acquisition had to be as short as possible. For this purpose, the present protocol focused on a minimal number of views (i.e., four echocardiographic views only), chosen to provide a wide range of quantitative variables for the assessment of heart dimensions and function using 2D and M-mode echocardiography, spectral Doppler, and STI, with all measurements performed after TTE sessions. Using the proposed method, a total of 28 variables of interest could be assessed offline, requiring a limited image acquisition duration, i.e., mean and maximal duration of respectively 3.8 and 6.3 minutes, only.

The apical 4- and 5-chamber views provided the largest number of quantitative variables. Interestingly, one single video loop of the apical 4-chamber view allowed the post-processing assessment of 12 2D variables (LA and LV diameters and volumes, LV systolic indices, mitral valve leaflets thicknesses) as well as one STI longitudinal LV StS variable (StS_4chv_). Similarly, one single video loop of the apical 5-chamber view allowed the post-processing assessment of three anatomic M-mode variables (LA and aortic diameters, LA:aorta ratio) as well as one STI longitudinal LV StS variable (StS_5chv_), with then calculation of the global LV StS (as the mean of StS_5chv_ and StS_4chv_). To summarize, a total of 18 variables could be assessed using only two video loops of apical views. The other 10 of the 28 tested imaging variables were technically more challenging because they needed the pre-processing placement of either the M-mode cursor (for mitral annular motion, TAPSE and indexed TAPSE measurements) or the sample volume and Doppler cursor for spectral Doppler variables (regarding diastolic transmitral flow and systolic arterial flows). Nevertheless, all within-day and between-day CV values of M-mode variables were <10%, and all those of spectral Doppler variables were <15%, with only two between-day CV values between 10 and 15% (12.1% and 13.4% for mitral A wave and mitral E:A ratio, respectively), which corresponds to good to excellent repeatability and reproducibility.

The present study also shows that all within-day and between-day CV values of STI variables were <10%. This is of particular interest as myocardial diseases, mainly including fibrosing cardiomyopathy and myocarditis, represent the most common orangutan heart diseases [1,4–6]. The ability of STI to early detect systolic myocardial dysfunction has been demonstrated in human patients with various subclinical myocardial diseases characterized by normal conventional echocardiographic parameters (including LV diameters, fractional shortening, and ejection fraction) [17,18]. We can therefore hypothesize that STI combined with spectral Doppler parameters of diastolic function could be useful for the early detection of myocardial diseases in captive BO.

In the present report, a significant BO effect was observed for all TTE variables, except for the LV fractional shortening. This result may be explained by the wide range of age (12 to 51 years old) and body weight (47 to 74 kg) of the four involved BO. As already described in the GAPH report [8], because orangutans vary in body shape and size, the optimal acoustic windows are slightly different from one animal to another. Additionally, among the four BO involved in the present study, the oldest female had the most prominent breast, making the transducer placement and images acquisition less easy. Finally, the four BO had variably developed laryngeal air sacs. These anatomical structures, typically extending along the ventral neck, beneath the clavicles and into the axilla, are involved in BO vocalization and communication [1]. Depending on their size, they may interfere with the ultrasound beam and variably alter the TTE image quality [8].

This preliminary work on TTE in BO presents several limitations. Quantitative measurements were limited to the left heart and aorta, as these were the easiest to detect and analyze. The right heart size was not evaluated, and TAPSE (indexed or not to body weight) was the only right ventricular function tested parameter. Additionally, our previous studies in dogs and cats showed that TTE is a highly observer-dependent examination [19,20]. The results presented here are thus only valid for the observer involved, and the authors encourage veterinarians to determine their own TTE variability, before undertaking further echocardiographic studies in BO or other great apes.

## Conclusions

In conclusion, our results demonstrate that TTE can be performed in awake BO after three months of animal training, providing the assessment of more than 20 2D, M-mode, spectral Doppler, and STI variables with good to excellent repeatability and reproducibility as well as minimal animal restraint. The non-stressful nature of the present method relies on several factors including the use of positive reinforcement for BO training to the technique, and rapidity of image acquisition (mean duration of 4 minutes), as only four 2D views are required, with subsequent offline post-processing analysis. This TTE method may be useful for longitudinal follow-up of heart morphology and function in awake BO, which is of particular interest as cardiovascular diseases have been identified as a major cause of mortality and morbidity in this endangered species.

## Acknowledgements

The authors would like to thank the Ménagerie, le Zoo du Jardin des Plantes staff for their collaboration in this study.

